# Exercise Leverages a Post-Stress Therapeutic Window in Hypothalamic CRH Neurons to Enable Recovery from Threat Sensitization

**DOI:** 10.64898/2026.01.07.698178

**Authors:** Mijail Rojas-Carvajal, Tamás Füzesi, Dinara Baimoukhametova, Nuria Daviu, Sarah Gibson Cook, Matthew Hill, Jaideep Bains

## Abstract

Exercise provides a range of benefits for physical and mental health. It relieves stress, yet paradoxically, it recruits the body’s stress response system. Here we show that activation of corticotropin-releasing hormone cells of the paraventricular nucleus of the hypothalamus (CRH^PVN^) and the endocrine arm of the stress axis is necessary, but not sufficient for the beneficial effects of exercise. Specifically, glucocorticoids act synergistically with brain-derived neurotrophic factor (BDNF) on CRH^PVN^ cells during exercise to reverse behavioral and synaptic sensitization after stress. In the absence of exercise, optogenetic activation of the tropomyosin-related kinase B (TrkB) receptor after stress is sufficient to reverse behavioral sensitization and synaptic metaplasticity. Our findings reveal a novel, time-sensitive mechanism by which exercise alleviates the impact of acute stress.

---

Exercise is widely recognized as a preventive strategy against various metabolic and neuropsychiatric conditions (1-3). Growing evidence supports the role of regular exercise in fostering stress resilience (4–6). Exercise also promotes adaptive responses to acute stress (4, 7, 8) and single bouts of exercise have immediate mood-enhancing effects (9). Paradoxically, exercise itself activates the hypothalamic pituitary adrenal (HPA) axis and increases circulating stress hormones (10–13). How exercise can elicit both a short-term stress response and engage compensatory neurobiological processes promoting long-term stress regulation is not known.

Here, we leveraged synaptic and behavioral readouts of stress sensitization to understand how exercise modulates the lasting consequences of acute stress. Using a combination of *ex vivo* electrophysiology, genetic, optogenetic, and behavioral approaches, we identify a critical period immediately after stress that may serve as a therapeutic window for the effects of exercise. Specifically, we show that exercise after stress increases both circulating corticosterone (CORT) and hypothalamic brain-derived neurotrophic factor (BDNF) concentrations. The synergistic action of these signaling elements in CRH^PVN^ neurons reverses behavioral threat sensitization and synaptic metaplasticity induced by stress. These findings provide the first mechanistic insights into how exercise alleviates the lasting effects of stress and reconcile previous observations regarding the timing and efficacy of exercise as a stress intervention.

## Results

### Exercise prolongs CRH^PVN^ activation and CORT secretion after stress but reverses its synaptic consequences when delivered within a critical period

Exercise relieves stress but also promotes endocrine stress signaling, highlighting a physiological paradox (12, 14). To understand this, we first assessed the effects of acute stress and exercise on two components of the HPA axis: the activity of CRH^PVN^ neurons and circulating concentrations of CORT. Mice expressing GCaMP6f in CRH^PVN^ neurons were unilaterally implanted with a ferrule for single fiber photometry (Fig. 1A-C). Baseline measurements were obtained in the home cage (HC). One group of mice received 10 foot shocks (FS) in a 5-min period; another group was placed on a treadmill and allowed to run for 1 hour (Fig. 1E). FS evoked a robust increase in CRH^PVN^ activity that returned to baseline levels ∼10 minutes after returning mice to HC (Fig. 1e, left panel). Exercise also evoked a robust increase in CRH^PVN^ activity that remained elevated through the entirety of running session (Fig. 1E, right panel). Tail blood was collected before experimental procedures started for quantifying baseline (BL) CORT levels. Then, a second sample was collected after treatments for post-stimuli measures (Post). This is consistent with the classical view that both acute stress and exercise increase CORT (Fig. 1F) and provides new information that exercise directly increases the activity of CRH^PVN^ neurons.

**Fig. 1.**
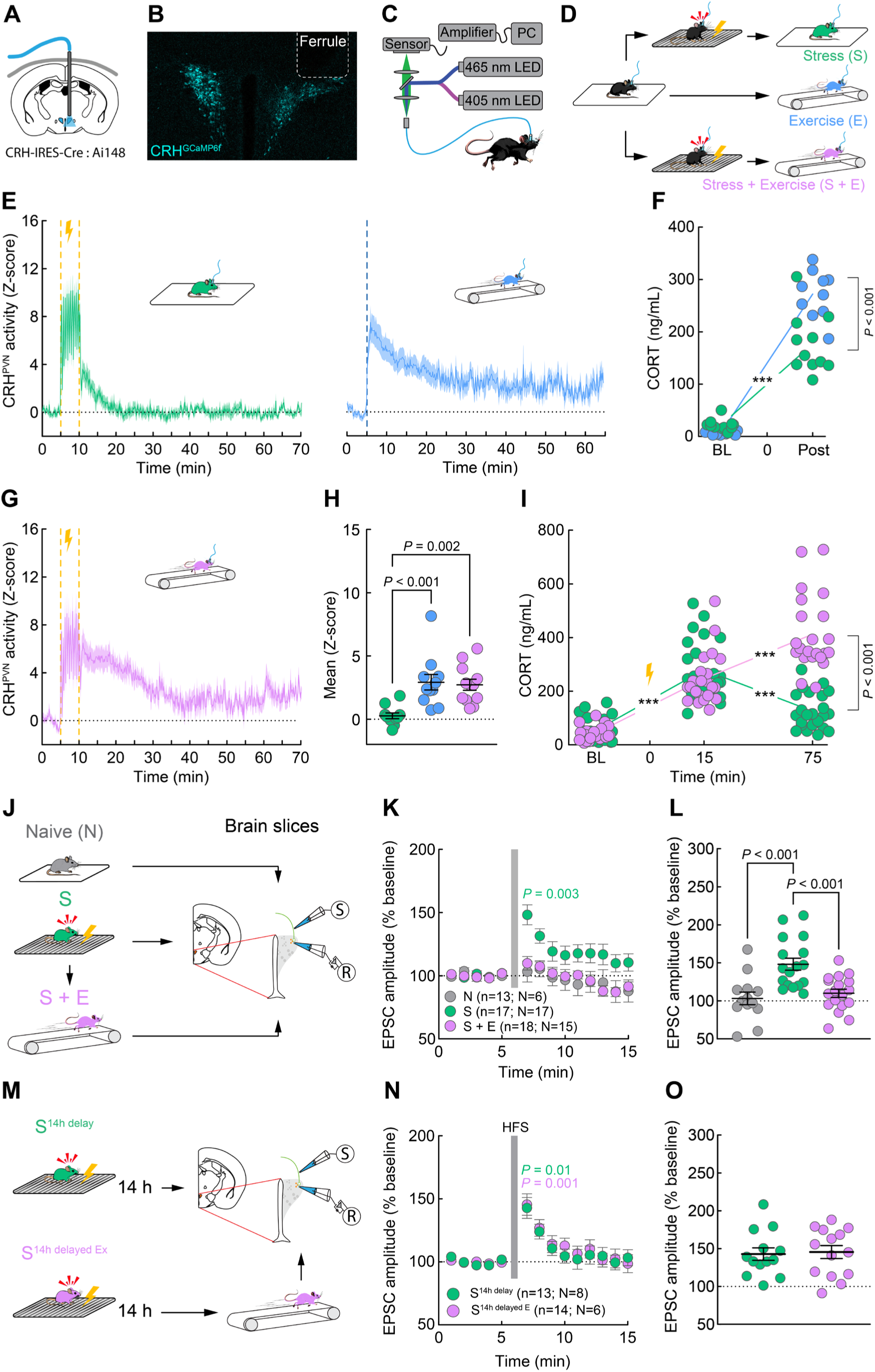
Exercise prolongs CRH activation and CORT after stress but reverses the synaptic consequences of stress when delivered within a sensitive window. **A**. Ferrule implantation for fiber photometry recordings. **B**. Representative confocal image of CRH^PVN^ neurons expressing GCaMP6f. **C**. Schematic of single-fiber photometry method. **D**. Experimental design. **E**. *In vivo* recordings of CRH^PVN^ activity in stress (S: N = 10), exercise (E: N = 11), and stress followed by exercise (S + E: N = 12) mice. **F**. CORT concentration from S (N = 11) and E mice (N = 10) before (baseline: BL) and after treatments (Post). Mixed ANOVA: Time (*F*(1, 19) = 330.1, *P* < 0.001), Group (*F*(1, 19) = 12.88, *P* = 0.002), and Time × Group (*F*(1, 19) = 22.63, *P* < 0.001). **G**. CRH^PVN^ activity in S+E mice. **H**. Average CRH^PVN^ activity after FS and during exercise. One-way ANOVA: *F*(2, 31) = 10.44, *P* < 0.001. **I**. CORT concentrations for S (N = 22) and S + E (N = 23) mice. Mixed ANOVA: Time (*F*(1.86, 80.00) = 136.2, *P* < 0.001), Group (*F*(1, 43) = 16.21, *P* < 0.001), and Time × Group (*F*(2, 86) = 68.43, *P* < 0.001). **J**. Whole-cell patch clamp recordings from CRH^PVN^ neurons in naïve (N), S, and S+E mice. **K**. EPSC amplitudes following HFS (grey bar) relative to BL (doted line). Mixed ANOVA: Time (*F*(4.434, 199.5) = 7.454, *P* < 0.001), Group (*F*(2, 45) = 7.682, *P* = 0.001), and Time × Group (*F*(8.867, 1995) = 3.232, *P* = 0.001). EPSC amplitude increased after HFS (min 5 vs min 7: *P* = 0.003) in S mice only. **L**. Average EPSC amplitude one minute after HFS. One-way ANOVA: *F*(2, 45) = 11.62, *P* < 0.001. **M**. Experimental design of whole-cell patch clamp **N**. EPSC amplitudes following HFS. Mixed ANOVA: Time (*F*(4.219, 105.5) = 16.44, *P* < 0.001). Both S^14h delay^ (*P* = 0.013) and S^14h delayed E^ (*P* = 0.005) mice showed increased EPSC amplitude after HFS. **O**. Average EPSC amplitude one minute after HFS was equally elevated in both groups. Data are mean ± SEM. ***: Within-group, *P* < 0.001.

Exercise is often used as a tool for stress relief. To better understand the connection between stress and exercise, we examined the effects of acute stress and exercise on CRH^PVN^ activity and CORT. Mice were subjected to FS and then placed on the treadmill immediately (Fig. 1G). As expected, FS strongly increased CRH^PVN^ activity. Exercise after stress sustained this increase in activity (Fig. 1G). When averaging CRH^PVN^ activity across the first hour after FS, mice that exercised after stress showed significantly higher activity levels compared to stressed mice that did not exercise (Fig. 1H). In contrast, FS did not alter overall CRH^PVN^ activity during running itself, as both exercise groups displayed comparable levels (Fig. 1H). To measure the effects of these treatments on CORT levels, we collected blood samples before stress, after FS, and after exercise. In FS mice, in the absence of exercise, CORT levels were increased 15 minutes after FS, then declined one hour later (Fig. 1I). In contrast, post-stress exercise further elevated CORT concentrations beyond those induced by stress alone (Fig. 1I). Both stress and exercise strongly activate the HPA axis, with post-stress exercise maintaining the elevation in CRH^PVN^ activity and slowing the recovery of the CORT response.

Next, we probed the effects of exercise after stress on a synaptic biomarker that has been identified after acute stress and may contribute to sensitization of this system: metaplasticity at glutamate synapses on CRH^PVN^ neurons (15, 16, 17). Following acute stress, glutamatergic synapses exhibit activity-dependent short-term potentiation (STP) that is not evident in naïve (unstressed) mice or rats. STP requires local CRH release (16) and can be elicited by optogenetic activation of CRH^PVN^ neurons in the absence of actual stress (17). To test the effects of exercise on STP after FS, we prepared brain slices from naïve and stressed mice, and from mice that exercised after stress, and whole-cell patch clamp recordings were obtained from CRH^PVN^ neurons (Fig. 1J). In naïve animals, high frequency stimulation (HFS) of glutamate inputs had no effect on excitatory postsynaptic current (EPSC) amplitude (Fig. 1K), consistent with previous reports that STP is not evident if animals are not stressed (15, 16, 17). As expected, STP was evident at glutamate synapses from stressed mice. We have previously shown that STP can be reliably induced up to three days after a single acute stress (15). If mice were subjected to exercise after FS, however, we observed no STP (Fig. 1L). This suggests that exercise, despite increasing CRH^PVN^ activity, attenuates STP after stress (Supplemental information Fig. S1). These observations show that although exercise causes a sustained increase in the activity of CRH^PVN^ neurons and amplifies HPA axis activation beyond stress levels, it counteracts the effects of stress on excitatory inputs onto CRH^PVN^ neurons.

Delivering interventions within a post-stress sensitive window may be critical to modulate synaptic plasticity such as STP (17, 18). Similar plasticity windows have been described (19), and evidence suggests that timely interventions can buffer the consequences of stress (20). To test whether delaying exercise after stress affected its capacity to buffer STP, mice subjected to FS were returned to HC (Fig. 1m) for fourteen hours before having access to exercise (S^14h delayed EX^) or remaining in their HC (S^14h delay^). Robust STP was detected in neurons from the control group S^14h delay^ mice (Fig. 1N), confirming the lasting impact of stress in CRH^PVN^ neurons (17). The S^14h delayed EX^ mice also showed a robust increase in EPSC amplitude that was indistinguishable from the S^14h delay^ mice (Fig. 1O). This evidence demonstrates that delayed exercise is ineffective in reversing the synaptic changes induced by acute stress, hinting at a potential sensitive window immediately after stress.

### Exercise reverses stress-induced behavioral threat sensitization

A single acute stress also induces behavioral sensitization (21–23). To probe the links between acute stress, exercise and behavioral threat response, we used the Dark-Light test (DL) which takes advantage of the fact that mice naturally avoid open, brightly illuminated spaces (24). We first investigated the effects of FS on multiple parameters in this test. Mice were habituated to the dark compartment and then placed in one of two groups: unstressed (FS context, no FS) and acute stress (FS) (Fig. 2A). The following day, mice were placed into the dark compartment of the DL chamber, and the connecting door was opened, providing free access to the light compartment for 5 minutes.

**Fig. 2.**
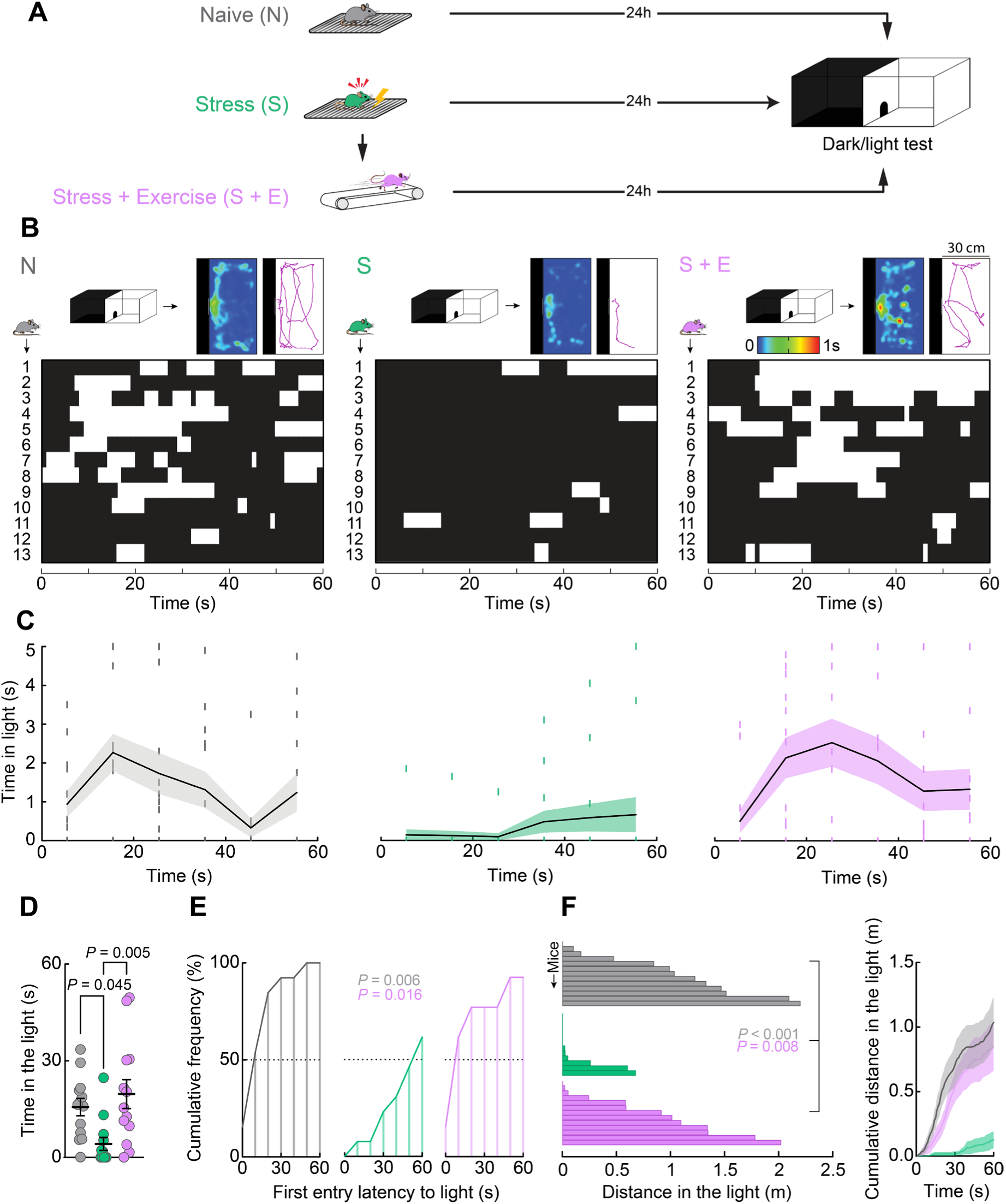
Exercise reverses stress-induced behavioral threat sensitization. **A**. Experimental design for the dark/light test (DL). Each group consisted of 13 male mice. Animals were tested in the DL 24 hours after treatment. **B**. Heat maps representing the spatial distribution of the average exploration time in the light compartment (top section of each panel). Tracking plot of the exploration trajectory of a representative animal per group. Behavioral time budget plot showing individual visits to the light (white bars) and dark compartment (black bars) of the DL test during the first minute of testing (bottom section of each panel). **C**. Frequency analysis of the exploration time in the light compartment binned every five seconds. **D**. Summary of time spent in the light compartment. One-way ANOVA: *F*(2, 36) = 6.132, *P* =.005). **E**. Frequency analysis of light compartment exploration latencies during the first minute of testing. One-way ANOVA: *F*(2, 18)= 7.569, *P*= 0.004. **F**. Distance travelled in the light compartment. One-way ANOVA: *F*(2, 36) = 8.900, *P* = 0.007). S showed a slow increase of cumulative distance over the first minute of test, with S + E and N showing a similar pattern of fast and steady increase (right panel). Tukey’s multiple comparisons test was used. Data are mean ± SEM.

In the control group, each mice entered the light compartment within the first minute, so we focused our analysis on this initial period (Supplemental information Fig. S2 for the complete 5-minute analysis). Analysis of movement in the light compartment revealed that naïve mice entered the compartment within the first minute (time to 1^st^ entry: 14.68 ± 3.761 s) and explored the entire area, with bouts of exploration interspersed with periods in the dark compartment (Fig. 2B, C, left panels). In contrast, stress exposure prolonged time to first entry into the light compartment, and suppressed exploration of the light compartment (Fig. 2B-F). Specifically, stressed mice showed a delay in the first exploration bout compared to naive mice (time to 1^st^ entry: 85.46 ± 17.46 s, *P* < 0.001 vs unstressed), with only 46% of the animals entering the light compartment within the first minute of testing (Fig. 2E; Supplemental information Fig. S2 G). Additionally, stressed mice covered significantly less distance while exploring the light compartment (Fig. 2F). The third group was subjected to FS, followed immediately by exercise. A single session of exercise fully reversed this behavioral phenotype (Fig. 2B, C). In mice subjected to stress followed by exercise, the time spent in the light compartment was higher than in stressed mice and comparable to that of naïve mice (Fig. 2D). Moreover, exercise restored the early exploration pattern observed in non-stressed mice (time to 1^st^ entry: 21.12 ± 5.65 s, *P* < 0.001 vs stressed), with 93% of these mice entering the light compartment within the first testing minute (Fig. 2E). Mice subjected to exercise after stress also traveled a greater distance than stressed mice, with cumulative levels over time being indistinguishable from those of naive mice (Fig. 2F). These findings demonstrate that a single session of exercise reverses stress-induced alterations in approach-avoidance dynamics, consistent with a decrease in behavioral sensitization to potential threat.

### Exercise has no effect on contextual fear memory

Aversive stimuli such as FS generate lasting conditioned fear memory, expressed as freezing when animals are re-exposed to the threat-associated context (25, 26). Since exercise following stress negates stress-induced metaplasticity and threat behavioral sensitization, we investigated whether it may impact the memory of the threat. Mice received FS and were either returned to HC or allowed to run on the treadmill (Fig. 3A). An additional group of naïve controls was exposed to the FS chamber for 5 minutes without receiving shocks. Twenty-four hours later, mice were re-exposed to the FS context for 5 minutes. Compared to naïve controls, FS context exposure elicited robust freezing in both stressed mice and those that exercised after stress, with no detectable differences between the stress and stress + exercise groups (Fig. 3B). Exploratory behaviors, including walking and rearing, were markedly reduced in these groups relative to naïve mice (Fig. 3C–D). Additionally, FS context elevated circulating CORT levels in both stress mice and mice exercising after stress, beyond those observed in naïve subjects (Fig. 3E). These observations indicate that exercise had no effect on the contextual fear memory or the CORT response.

**Fig. 3.**
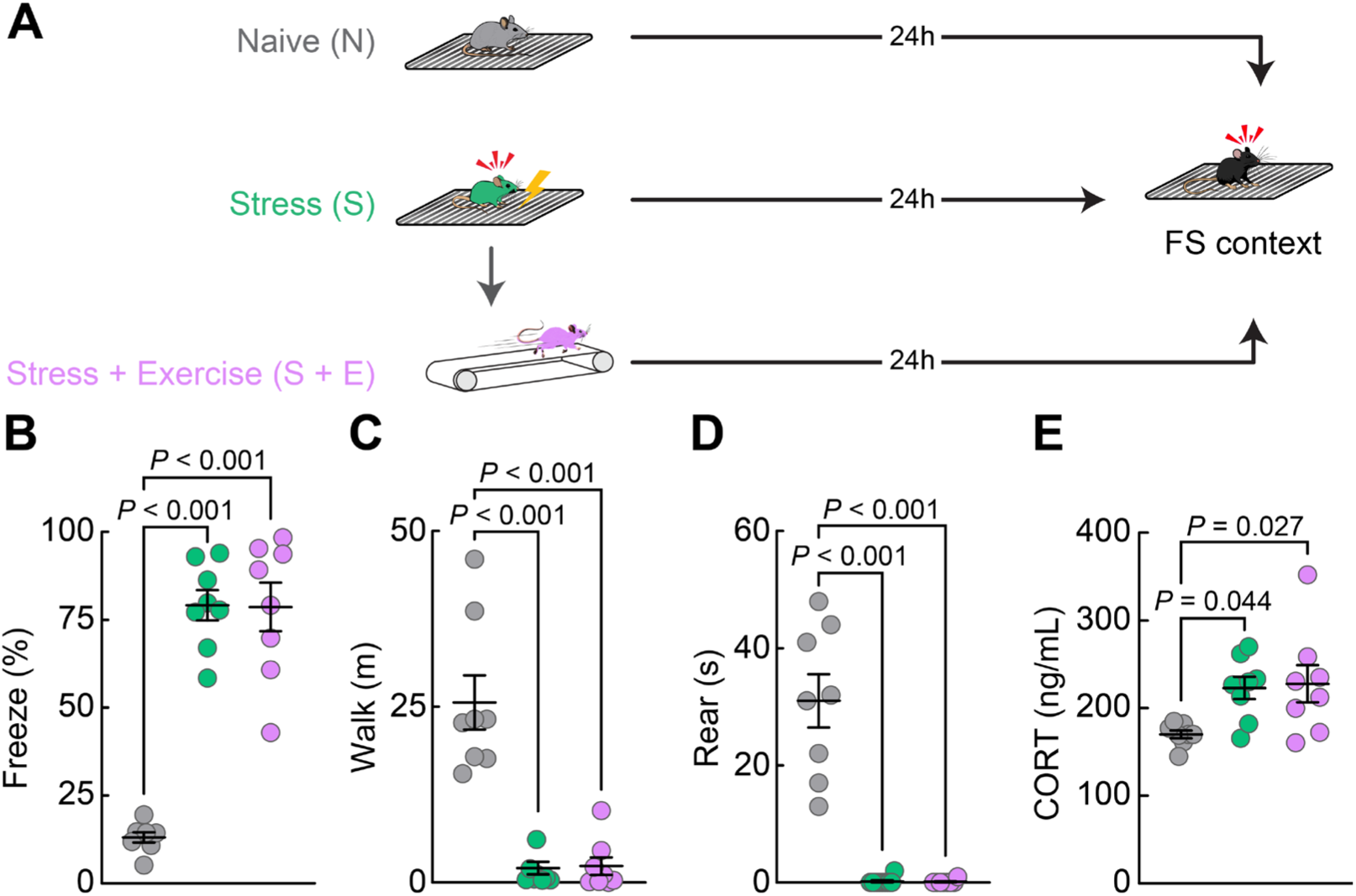
Exercise has no effect on contextual fear memory. **A**. Experimental design for contextual fear conditioning. Each group consisted of 8 male mice. Mice in all groups were exposed to the FS chamber for 5 minutes. Stress (S) and Stress + Exercise (S+E) mice were exposed to FS and then either returned to their HC (S) or placed on a treadmill for 1 hour of running (S+E). Naïve (N) mice were not exposed to FS and were returned directly to HC. Twenty-four hours later, mice were re-exposed to the FS chamber for 5 minutes and the following behaviors were analyzed automatically: freezing (percentage of total time), walking (meters), and rearing (seconds). Mice returned to HC after that, and tail blood was collected 15 minutes later for CORT quantification using ELISAS. **B**. Individual freezing percentage per group. One-way ANOVA: *F*(2, 21) = 63.13, *P* < 0.001. **C**. Individual distance travelled per group. One-way ANOVA: *F*(2, 21) = 33.32, *P* < 0.001. **D**. Individual rearing duration. One-way ANOVA: *F*(2, 21) = 34.37, *P* < 0.001. **E**. Individual CORT concentration per group. One-way ANOVA: *F*(2,21) = 4.90, *P* = 0.018. Tukey’s multiple comparisons test was used. Data are mean ± SEM.

### Elevated CORT synergizes with a TrkB ligand to prevent stress-induced metaplasticity

Post-stress exercise elevates HPA axis activity beyond stress levels, yet it buffers the synaptic consequences of stress. One possible explanation is that the prolonged elevation in CORT is sufficient to buffer the effects of stress. To test whether the increased and prolonged CORT output is sufficient to deter STP expression, we mimicked exercise-induced modulation of the HPA axis by using a 1-hour immobilization (IMMO) protocol. Mice were unilaterally implanted for fiber photometry recordings of CRH^PVN^ activity during IMMO (Fig. 4A). IMMO increased CRH^PVN^ activity for the duration of the session (Fig. 4A). As expected, CORT was markedly elevated 1 hour after IMMO onset (Fig. 4B). Next, we evaluated the effects of IMMO on STP (Fig. 4C, top section). Whole-cell patch clamp recordings revealed robust STP (Fig.4C, bottom section). Thus, prolonged CRH^PVN^ activity and elevated CORT do not deter the expression of STP. It is unclear, however, whether CORT itself after stress would affect STP. To test this idea, brain slices from stressed mice were incubated in a bath containing artificial cerebrospinal fluid (aCSF; S^aCSF^) or CORT (100 nM) diluted in aCSF (S^CORT^) for 1 hour (Fig. 4D). Whole-cell patch clamp recordings of both S^aCSF^ and S^CORT^ mice showed STP (Fig. 4e). Together, these observations indicate that the elevated circulating CORT induced by post-stress exercise is insufficient to prevent STP, suggesting that a distinct molecular mediator may be responsible for the buffering effects of exercise.

**Fig. 4.**
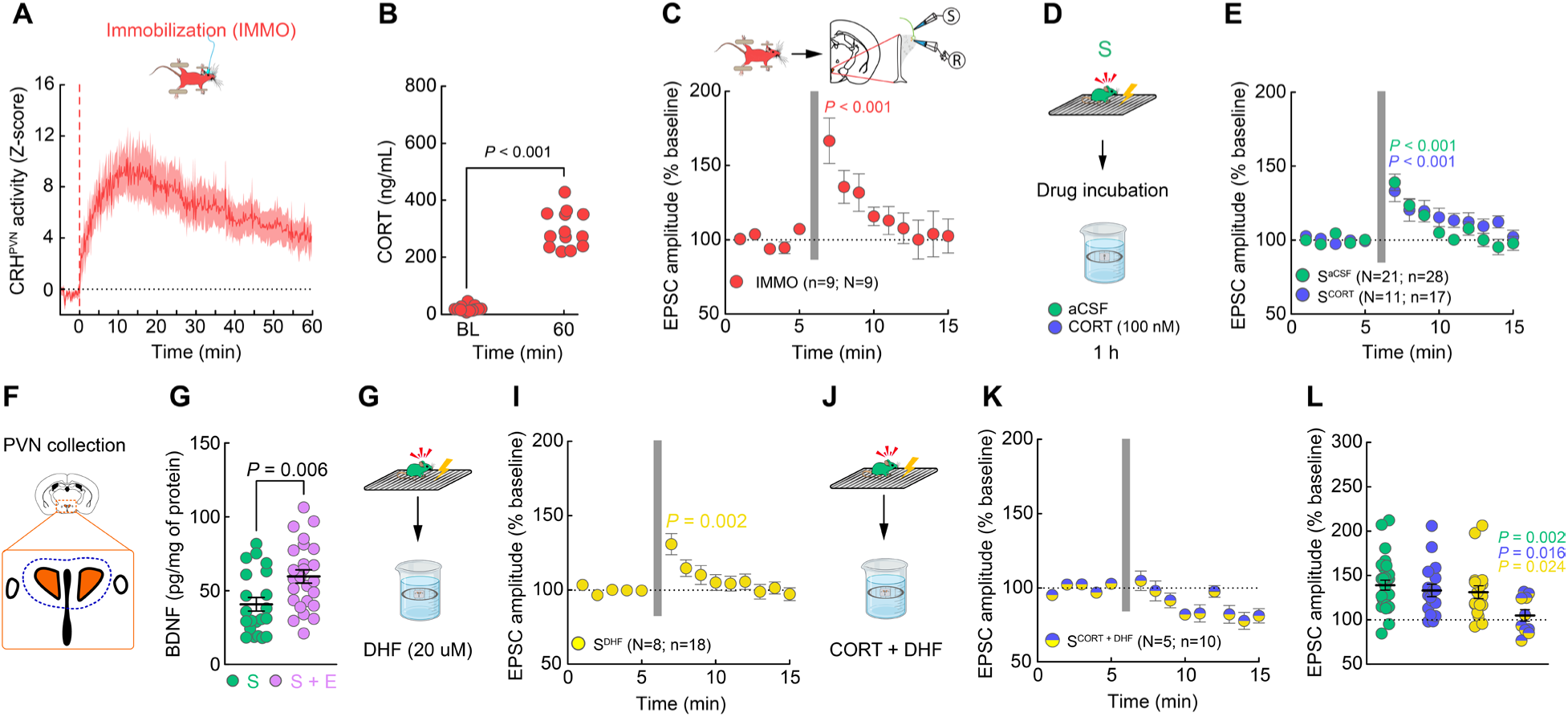
Elevated CORT synergizes with a TrkB ligand to prevent stress-induced metaplasticity. **A**. CRH^PVN^ calcium response (Z-score) during one hour of immobilization (IMMO; N = 9). IMMO elevated average CRH^PVN^ calcium response compared to BL (two-tailed, one sample *t*-test: *t*(7)=4.478, *P* = 0.003). **B**. Quantification of plasma concentrations of CORT (N = 13) before and after IMMO. Two-tailed paired *t*-test: *t*(12)=15.92, *P* < 0.001. **C**. Schematic of whole-cell patch clamp recordings from CRH^PVN^ neurons after 1-hour IMMO (top section). Repeated-measures ANOVA: *F* (13, 104) = 6.762, *P* < 0.001; min 5 vs min 7: *P* < 0.001. **D**. Experimental design for CORT incubation of brain slices. **E**. A mixed ANOVA revealed a main effect of Time (*F* (5.485, 234.6) = 16.85, *P* < 0.001). **F**. Schematic of PVN tissue collection for protein quantification of brain-derived neurotrophic factor (BDNF). **G**. Individual concentration of BDNF in the PVN. Two tail *t*-test: *t*(44)=2.901, *P* = 0.006. **H**. Experimental design for 7,8-Dihydroxyflavone (DHF) incubation of brain slices. **I**. CRH^PVN^ neurons from S^DHF^ mice showed a significant increase in EPSC amplitude after HFS (repeated measures ANOVA: *F* (13, 221) = 5.791, *P* < 0.001; min 5 vs min 7: *P* < 0.001). **J**. Experimental design for CORT + DHF incubation of brain slices. **K**. Following HFS, no changes were observed in the EPSC amplitudes of CRH^PVN^ neurons in S^CORT + DHF^ mice. **L**. Summary of the average EPSC amplitude per cell in all drug-treated slices. One-way ANOVA: *F* (3, 69) = 3.490, *P* = 0.020. Tukey’s and Fisher’s LSD multiple comparisons tests were used. Geisser-Greenhouse correction was used. Data are mean ± SEM.

Regular exercise promotes BDNF release, a neurotrophin critical for neuronal growth, survival, and diverse forms of synaptic plasticity, which acts primarily through its high-affinity receptor, tropomyosin receptor kinase B (TrkB) (27–32). The capacity of a single exercise bout to elevate BDNF varies between brain regions (33, 34), and to our knowledge, it is unknown whether acute exercise elevates BDNF in the PVN, especially in stressed mice. Therefore, we assessed BDNF concentrations in bilateral PVN samples from stressed mice and mice that exercised after stress (Fig. 4F). Protein quantification revealed a significant increase in BDNF concentrations in mice that exercised after stress compared to stressed controls (Fig. 4G). Since post-stress exercise locally increases BDNF in the PVN, we next tested whether activating TrkB receptors in the PVN of stressed mice could buffer STP. Brain slices were incubated for 1 hour in aCSF containing the TrkB ligand 7,8-Dihydroxyflavone (DHF; 20 µM; S^DHF^ condition; Fig. 4H) (35–37). Whole-cell patch clamp recordings showed STP after HFS in the S^DHF^ group (Fig. 4I), indicating that TrkB activation after stress is insufficient to prevent STP.

Glucocorticoid receptors (GR) and TrkB share key intracellular signaling pathways (38), and some forms of synaptic plasticity require a synergistic interaction between CORT and BDNF (39). We hypothesized that post-stress exercise may induce adaptive GR/TrkB activation in a time-dependent manner, contributing to STP buffering. To test this, brain slices from stressed mice were incubated for 1 hour in aCSF containing both CORT and DHF (S^CORT + DHF^ condition; Fig. 3J). STP was fully prevented in this group (Fig. 4K). EPSC amplitudes during the first minute post-HFS were significantly reduced in S^CORT + DHF^ cells compared to S^aCSF^, S^CORT^, and S^DHF^ conditions (Fig. 4L). These findings indicate that the co-activation of GR and TrkB is sufficient to fully block STP *in vitro*.

### Post-stress activation of TrkB in CRH^PVN^ cells prevents stress-induced STP and anxiety-like behaviors

Next, we asked whether the effects of TrkB activation and CORT on CRH^PVN^ neurons were necessary and sufficient to reverse the effects of stress. To address this question, we selectively disrupted TrkB signaling by overexpressing a truncated version of this receptor (TrkB.T1) in CRH^PVN^ neurons, then tested whether post-stress exercise (S + E^TrkB.T1^) was still capable of buffering STP (Fig. 5A-C). CRH^PVN^ neurons from S + E^TrkB.T1^ mice showed robust STP (Fig. 4D), demonstrating that TrkB activation in CRH^PVN^ neurons during exercise is necessary to buffer the synaptic effects of stress. Importantly, overexpression of TrkB.T1 in CRH^PVN^ neurons did not induce STP or block stress-induced STP (Supplemental information Fig. S3 A-B). DHF buffered STP in the presence of CORT, but this does not indicate whether direct TrkB activation in CRH^PVN^ neurons after stress –but in the absence of exercise– is sufficient to buffer STP. We tested this idea by expressing a light-activated TrkB (Opto-cytTrkB) (40) in CRH^PVN^ neurons and delivering light stimulation through an optic fiber for 15 minutes immediately after FS (S^Opto-TrkB^; Fig. 4A-C). Since circulating CORT is rapidly elevated after FS (Fig. 1), no exogenous CORT was supplemented. We observed no STP in CRH^PVN^ neurons of S^Opto-TrkB^ mice (Fig. 5E; Supplemental information Fig. S3 D). During the first minute post-HFS, EPSC amplitudes were smaller in S^Opto-TrkB^ compared to S + E^TrkB.T1^ mice (Fig. 5F). This demonstrates that, in the presence of elevated CORT, TrkB activation in CRH^PVN^ cells is necessary and sufficient to buffer stress-induced STP.

**Fig. 5.**
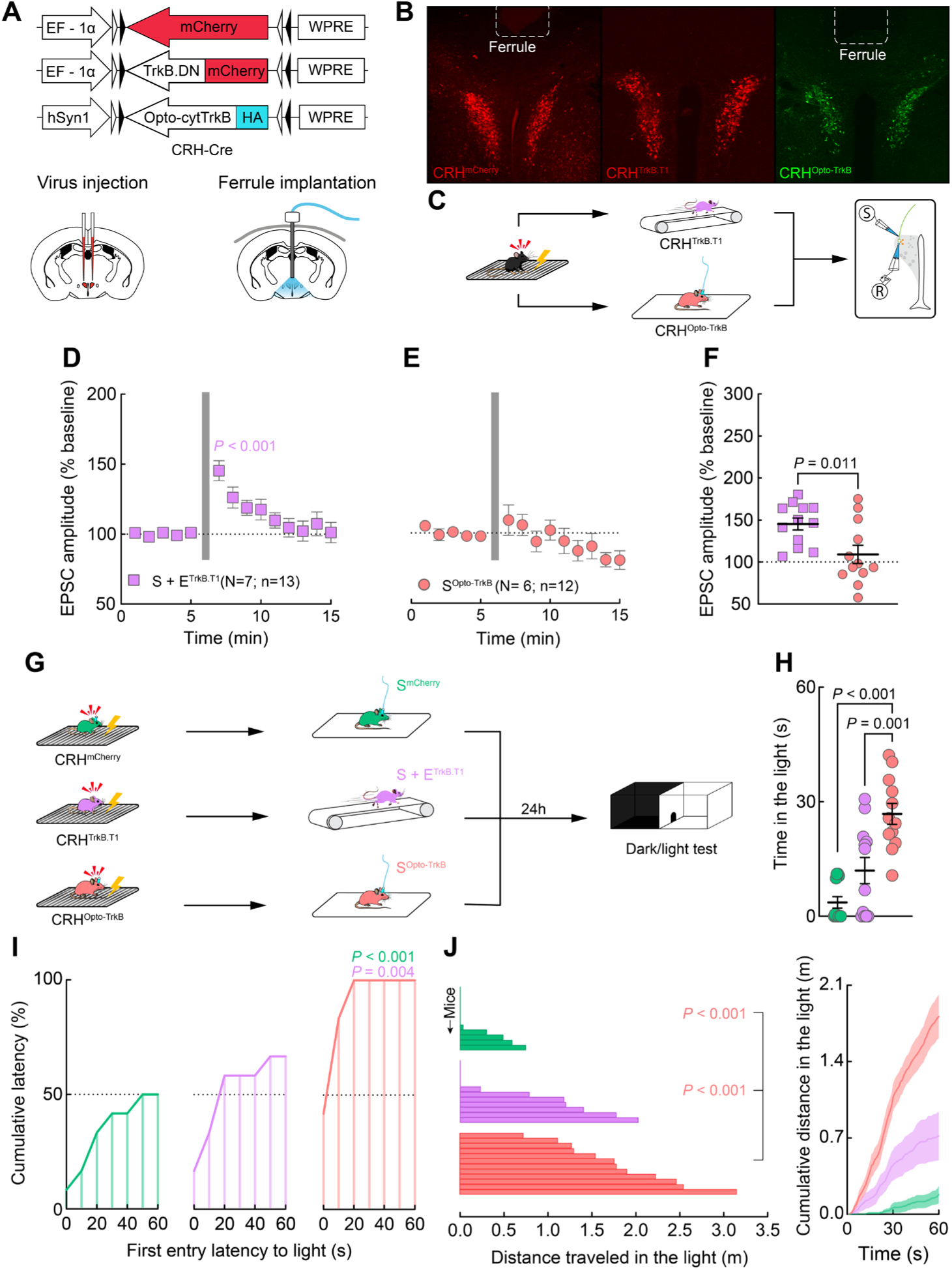
Post-stress activation of TrkB in CRH^PVN^ cells reverses stress-induced STP and threat behavioral sensitization. **A.** Constructs of the following Cre-dependent viruses: AAV1-hSyn-DIO-mCherry, AAV1-EF1a-DIO-trkB.DN-mCherry, and AAV1-hSyn1-DIO-Opto-cytTrkB(E281A)-HA (top section). Schematic showing bilateral virus injection in the PVN (bottom left). Schematic of ferrule implantation for optogenetic stimulation of the PVN (bottom right). **B**. Confirmatory confocal image of virus expression in CRH^PVN^ cells. Traces of the ferrule implantation in the tissue are delineated. **C**. Experimental design for genetic and optogenetic manipulation of TrkB after stress. **D**. After HFS (grey bar), CRH^PVN^ neurons from S + E^TrkB.T1^ showed a significant increase in EPSC amplitudes (repeated measures ANOVA: *F* (13, 143) = 10.53, *P* < 0.001, min 5 vs min 7: *P* < 0.001). **E**. CRH^PVN^ neurons from S^OptoTrkB^ showed no changes in EPSC amplitudes after HFS. **F**. Average EPSC amplitude per cell one minute after HFS. Unpaired two-tailed t-test: *t*(22)=2.777, *P* = 0.011. **F**. Experimental design for genetic and optogenetic manipulation of TrkB after stress and its effects on the DL test. **H**. Time in the light compartment during the first minute of testing. One-way ANOVA: *F* (2, 33) = 19.13, *P* < 0.001). **I**. Frequency analysis of light compartment exploration latencies. The percentage of both S^mCherry^ (*P* < 0.001) and S + E^TrkB.T1^ mice (*P* < 0.004) initiating their first exploration bout in the light compartment was lower than in S^OptoTrkB^ mice (One-way ANOVA: *F* (2, 18) = 15.03, *P* < 0.001). **J**. Analysis of the distance travelled in the light compartment. Both S^mCherry^ (*P* < 0.001) and S + E^TrkB.T1^ mice (*P* < 0.001) travelled less distance than in S^OptoTrkB^ mice (One-way ANOVA: *F* (2, 33) = 21.57, *P* < 0.001). Cumulative distance travelled in the light compartment (right panel). Tukey’s multiple comparisons test and Geisser-Greenhouse correction were used. Data are mean ± SEM.

Next, we tested whether these molecular and electrophysiological changes translate into behavioral outcomes. Specifically, we asked if TrkB activation in CRH^PVN^ neurons is necessary and sufficient to reverse threat behavioral sensitization following stress. Thus, S + E^TrkB.T1^ and S^Opto-TrkB^ mice were tested in the DL box 24 hours after treatment (Fig. 5G). An additional group of mice expressing mCherry in CRH^PVN^ neurons was implanted with an optic fiber and light was delivered after FS (Fig. 4A, G), serving as a stress control group (S^mCherry^). The analysis revealed sparse and delayed light-compartment exploration bouts in S^mCherry^ mice, increased but delayed visits in S + E^TrkB.T1^ mice, while prompt, frequent exploration in S^Opto-TrkB^ mice (Supplemental information Fig. S4 A–B; Five-minutes analysis: Supplemental information Fig. S4 D-I). The analysis of visit durations revealed that S^mCherry^ mice spent most of their time in the dark compartment (Fig. 5H). Similarly, S + E^TrkB.T1^ mice spent limited time in the light, although their stay was descriptively longer compared to S^mCherry^ mice. In contrast, post-FS optical stimulation of TrkB fully reversed the effects of stress, with S^Opto-TrkB^ mice showing the longest duration in the light compartment compared to all other groups (Fig. 5H). The analysis of latencies confirmed delayed first exploration bouts in S^mCherry^ mice, with only 50% of them entering the light compartment within the first testing minute (Fig. 5I). In S + E^TrkB.T1^ mice, shorter latencies were detected, with only 70% of the animals vising the light compartment during the same period. In contrast, 100% of S^Opto-TrkB^ animals explored the light compartment within the first minute of testing, exhibiting the shortest exploration latencies among any of the groups (Fig. 5I). No significant difference was detected between S^mCherry^ and S + E^TrkB.T1^ animals. The analysis of distance traveled revealed a similar pattern. S^mCherry^ mice exhibited the shortest displacement within the light compartment, followed by S + E^TrkB.T1^ animals, whose increased exploration did not reach statistical significance compared to S^mCherry^ mice (Fig. 5J, left panel). In contrast, S^Opto-TrkB^ mice showed the most extensive exploration across all groups. Exploration kinetics further revealed that early light-compartment exploration in S^Opto-TrkB^ mice was accompanied by greater cumulative distances traveled, a pattern that was less pronounced in the other groups. (Fig. 5J, right panel). These findings demonstrate that TrkB signaling plays a fundamental role in mediating the behavioral effects of post-stress exercise. While S + E^TrkB.T1^ animals exhibited a partial recovery from the stress-induced threat behavioral sensitization, the expression of CRH^TrkB.T1^ attenuated the otherwise robust and consistent behavioral benefits of post-stress exercise. Moreover, the ability of optogenetic TrkB activation to fully reverse stress-induced behavioral sensitization highlights a critical role for TrkB signaling in CRH^PVN^ neurons in modulating the brain’s response to stress.

## Discussion

The ability of exercise to activate the HPA axis presents a physiological paradox, complicating efforts to delineate the neurobiological mechanisms by which it exerts stress-relieving effects. Here, we demonstrate that a single bout of aerobic exercise, immediately after acute stress, selectively reverses stress-induced threat sensitization and synaptic metaplasticity in CRH^PVN^ neurons, while sparing adaptive fear memory. This reversal requires a localized, exercise-induced increase in BDNF within the PVN, which when coinciding with elevated CORT, activates TrkB receptors in CRH^PVN^ neurons to promote adaptive plasticity. Notably, this effect occurs only when exercise is performed after stress exposure, revealing a sensitive window during which stress-induced synaptic changes remain liable and subject to intervention.

CORT and BDNF exhibit complex, temporally dynamic interactions in the PVN. Acute stress increases circulating CORT within 15 minutes while reducing BDNF protein content and simultaneously elevating BDNF mRNA expression in parvocellular PVN neurons (41–43). PVN BDNF content increases 1 hour after stress (41), and central BDNF administration increases circulating CORT within 30 minutes (42). Conversely, prolonged acute stress reduces BDNF mRNA in the PVN without affecting BDNF content (42). GR activity suppresses TrkB signaling but BDNF/TrkB activation does not alter GR function (43). In our study, both short and prolonged acute stress induced STP. Such plasticity was unaffected by activating GR or TrkB alone, but simultaneous activation of both pathways fully reversed STP, highlighting the necessity of temporal coordination and combinatorial signaling. Although stress can elevate both BDNF and CORT in the PVN, their endogenous release is either temporally misaligned or subthreshold, rendering them ineffective to prevent or reverse synaptic metaplasticity. This points to the necessity of synchronizing these signals post-stress to effectively modulate plasticity and mitigate stress-related circuit changes.

The capacity of acute exercise to increase BDNF in subcortical regions like the PVN is not well described. Treadmill running elevates BDNF content in sensorimotor cortex but not in the hippocampus (34). However, it increases hippocampal BDNF mRNA shortly after exercise and for several minutes (45, 46), with BDNF content increasing 6 hours later (47). Running also increases BDNF in the prefrontal cortex (48). Importantly, treadmill exercise sensitizes TrkB receptors in the hippocampus, promoting stronger and prolonged activation (50). In contrast, acute treadmill running reliably engages the HPA axis, inducing a dose-dependent activation of CRH^PVN^ cells (14, 50) and increasing circulating CORT (34, 47, 51, 52). Here, we demonstrated that acute post-stress treadmill exercise increases BDNF concentrations in the PVN and provided *in vivo* evidence of the temporal dynamics of CRH^PVN^ activity following exercise. We also demonstrated that the temporally coordinated rise in both BDNF and CORT levels within the PVN following exercise is both necessary and sufficient to reverse stress-induced STP in CRH^PVN^ neurons. Our findings suggest a hormone-dependent gating mechanism whereby glucocorticoids prime CRH^PVN^ neurons to respond adaptively to environmental interventions. The intracellular mechanisms by which CORT enables TrkB activation to buffers stress-induced plasticity within the post-stress critical window remain to be elucidated. STP expression in CRH^PVN^ neurons requires a downregulation of NMDA receptors (16). It is possible that enhanced BDNF/TrkB signaling post-exercise may promote increased intracellular Ca^2+^ release through canonical (29, 53, 54) and non-canonical pathways (55), preventing STP. Further investigation is required to determine the precise intracellular mechanism involved.

This work shows that, although different stimuli including exercise activate CRH^PVN^ neurons, only stressors prime them for STP. This indicates that BDNF may act as the molecular mediator shifting CRH^PVN^ activation from driving synaptic metaplasticity to enabling stress buffering. Notably, CORT negative feedback was insufficient to prevent STP. While CORT rapidly suppresses CRH^PVN^ neuronal activity (59), it did not block the synaptic metaplasticity, underscoring a dissociation between hormonal feedback and local synaptic plasticity mechanisms. Given the local effects of BDNF/TrkB signaling in CRH^PVN^ neurons on threat behavioral sensitization, our findings support that stress-induced synaptic metaplasticity serves as a cellular correlate of stress memory. The persistence of this metaplasticity days after stress (17) and its association with behavioral sensitization suggests that CRH^PVN^ neurons maintain a latent “readiness” that encodes prior aversive experience. This synaptic trace may act as a predictive internal model of future threat. Our observations also underscore the critical role of CRH^PVN^ neurons in integrating aversive and rewarding signals to guide behavior (17, 56–58). Interestingly, this phenomenon diverges from canonical contextual fear memory. Contextual fear serves an adaptive role by minimizing predator detection and promoting energy conservation (25, 26). Exercise selectively buffers CRH^PVN^ plasticity and its associated behaviors while leaving the adaptive processes of contextual fear learning that promote threat avoidance and survival intact.

Clinical evidence suggests that a single bout of exercise rapidly enhances mood (9, 60), reduces anxiety and depressive symptoms (62, 63), and attenuates HPA axis response to acute stress (4, 63). However, the mechanisms by which acute exercise modulates stress responsiveness remain understudied. Building upon these clinical insights, our preclinical work uncovered a possible “therapeutic” window during which lasting stress effects can be reversed. Given the role of CRH^PVN^ neurons in anxiety and stress (64–68), our findings provide a mechanistic basis for this intervention and highlight a novel, non-invasive strategy for remodeling stress-sensitive circuits. Importantly, the existence of this therapeutic window opens new avenues for pharmacological and non-pharmacological interventions. Although efforts to develop TrkB agonists (e.g., positive allosteric modulators) have faced challenges and remain in early stages, several existing compounds such as ketamine and certain psychedelics, engage BDNF/TrkB signaling pathways (69–71). This raises the exciting possibility that post-stress interventions – whether behavioral or pharmacological – could be optimized to target a temporally defined window of plasticity within stress-responsive brain regions. Together, these insights pave the way for the development of time-sensitive, mechanism-based strategies to prevent or reverse the maladaptive effects of acute stress.

## Materials and Methods

### Subjects and general procedures

Mice strains with a C57BL/6 background were used (Jackson Laboratory). Males and female mice were used in all experiments unless specified. For whole-cell patch-clamp recordings, a mouse line expressing Cre in a CRH-dependent manner (CRH-IRES-Cre) was crossed with a Cre-dependent tdTomato reporter line (Ai14) to generate mice with tdTomato-labeled CRH⁺ cells (CRH-IRES-Cre × Ai14) (*76*). For *in vivo* fiber photometry experiments, the Ai148 mouse line – which expresses the calcium sensor GCaMP6s in a Cre-dependent manner – was crossed with the CRH-IRES-Cre line to generate mice expressing GCaMP6s specifically in CRH⁺ cells (Ai148 × CRH-IRES-Cre) (66). Cre-dependent adeno-associated viruses (AAVs) were injected bilaterally into CRH-IRES-Cre or Ai148 × CRH-IRES-Cre mice, to obtain viral expression in CRH^+^ cells only. For behavioral studies, all mouse lines were used. For all *in vivo* experiments, animals were habituated to experimental handling using the following procedure: once daily for three consecutive days, mice were scooped and held in the experimenter’s hands for 1 minute, followed by 1-minute rest periods, for a total of 10 minutes per session under dimmed light. Mice used in fiber photometry experiments were additionally habituated to fiber optic attachment and detachment. This was accomplished by gradually increasing the level of restraint during the handling sessions. Over the final two days of habituation, mice were fitted with fiber optic cables for 20 minutes per day while freely exploring their home cages.

Mice were 6–8 weeks old at the time of surgical procedures and viral injections, and 8–10 weeks old at the time the experiments. All male mice were single housed for at least one week prior to the start of experimental procedures. To avoid sex-specific effects of acute social isolation or social buffering on PVN electrophysiology (Sterley et al., 2018), female mice were group-housed until the start of experimental procedures. Female mice used in *in vivo* experiments were single-housed one week prior to the beginning of testing. All animals were housed in transparent polycarbonate cages (35.5 × 17.5 × 13 cm) under a 12:12-hour light–dark cycle (lights on at 7:00 a.m.), with *ad libitum* access to food and water.

### Stereotaxic procedures

Mice were anesthetized with isoflurane during all surgical procedures. For in vivo fiber photometry recordings, a 400 µm-diameter mono fiber optic ferrule (Doric Lenses; MFC_400/430-0.48_5.5mm_MF2.5_FLT) was implanted targeting the PVN using the following stereotaxic coordinates: anterior–posterior (AP): −0.7 mm; medial–lateral (ML): ±0.5 mm; dorsal–ventral (DV): −4.8 mm from the dura. To minimize tissue damage, a stainless-steel tracking ferrule (diameter: 0.33 mm) was first lowered at a rate of 1.5 mm/min to a depth of DV: −4.5 mm, then immediately withdrawn. The optical ferrule was subsequently lowered to the final target depth at 0.2 mm/min. The ferrule was secured to the skull using Metabond and dental cement. For viral injections, a pulled glass micropipette containing the viral construct was lowered to the PVN (AP: −0.7 mm, ML:±0.5 mm, DV: −4.7 mm from the dura), and a total volume of 210 nL was delivered via pressure injection (Nanoject II, Drummond Scientific). After injection, the micropipette was left in place for 5 minutes to allow for viral diffusion before being slowly retracted. Postoperatively, mice received analgesia (Meloxicam, 5 mg/kg, subcutaneous) 24 hours after surgery and were monitored daily for 7 days. All mice were given a 2-week recovery period before experimental procedures started.

#### Viruses

The following Cre-dependent AAV plasmids (pAAV) were obtained from Addgene and packaged into AAV1 vectors: pAAV-hSyn1-DIO-Opto-cytTrkB(E281A)-HA (5.85 × 10¹³ GC/mL; Addgene #180590) and pAAV-EF1a-DIO-trkB.DN-mCherry (1.95 × 10¹³ GC/mL; Addgene #121502). The AAV1-hSyn1-DIO-Opto-cytTrkB(E281A)-HA was used to enable optogenetic manipulation of TrkB signaling, while AAV1-EF1a-DIO-trkB.DN-mCherry was used to reduce endogenous TrkB activity via expression of a dominant-negative mutant. Both viral vectors were stereotaxically injected into the PVN of CRH-IRES-Cre or Ai148 × CRH-IRES-Cre mice for cell-type-specific expression in CRH⁺ neurons.

### Histology

To verify viral expression and ferrule placement, mice were anesthetized with isoflurane and their brains rapidly extracted and submerged in ice-cold aCSF for approximately 5 minutes. Coronal brain slices (200 µm) containing the PVN were prepared using a vibratome (VT1200 S, Leica) in ice-cold aCSF. Slices were fixed in 4% paraformaldehyde (PFA) in phosphate buffer (PB) at 4 °C for 5 minutes, followed by immersion in 30% sucrose in phosphate-buffered saline (PBS) for an additional 5 minutes. For animals injected with HA-tagged viral constructs, immunohistochemistry was performed. Slices were incubated for 24 h at 4 °C with a primary antibody against the HA epitope (1:1000 dilution; rabbit monoclonal antibody, C29F4; Cell Signaling Technology). After three washes in PBS containing 0.3% Triton X-100, slices were incubated with a secondary Alexa Fluor 488-conjugated anti-rabbit antibody (1:1000 dilution; Cell Signaling Technology) for 2 h at room temperature. Following final washes in PBS, slices were mounted and coverslipped using Vectashield mounting medium. Images were acquired using a confocal microscope (Olympus BX50 Fluoview) and processed using ImageJ (NIH).

### Slice preparation and ex vivo electrophysiology

All preparations and recordings were done as previously reported (15; 27). Mice were anesthetized (Isoflurane) and decapitated 5-10 minutes after experimental manipulations end. The brain was removed and submerged into ice-cold slicing solution containing (in nM): 87 NaCl, 2.5 KCl, 0.5 CaCl_2_, 7 MgCl_2_, 25 NaHCO_3_, 25 d-glucose, 1.25 NaH_2_PO_4_ and 75 sucrose, saturated with 95% O_2_/5% CO_2_. Coronal slides (250 µm) obtained using a vibratome (VT 1200 S, Leica) were kept for 15 minutes in a 30°C N-methyl-d-glucamine (NMDG)-recovery solution containing (in nM): 2.5 KCI, 25 NaHCO_3_, 0.5 CaCl_2_, 10 MgCl_2_, 1.2 NaH_2_PO_4_, 25 glucose, 110 NMDG and 110 HCI, saturated with 95% O_2_/5% CO_2_. Then, slides were incubated for 1 hour in 30 °C artificial cerebrospinal fluid (aCSF) containing (in mM): 126 NaCl, 2.5 KCl, 26 NaHCO_3_, 2.5 CaCl_2_, 1.5 MgCl_2_, 1.25 NaH_2_PO_4_ and 10 glucose, saturated with 95% O_2_/5% CO_2_.

#### Electrophysiological recording protocol

All recordings were obtained in aCSF containing picrotoxin (100 μM) at 30-32 °C, perfused at 1 mL/min. Neurons were visualized using an upright microscope fitted with differential interference contrast and epifluorescence optics and a camera. Borosilicate pipettes (2.5 - 4.5 mΩ) were filled with internal solution containing (in mM): 108 K-gluconate, 2 MgCl_2_, 8 sodium-gluconate, 8 KCl, 1 K-EGTA, 4 K-ATP, 0.3 Na-GTP and 10 HEPES buffer. Current-clamp recordings were acquired at a −70 mV membrane potential. To assess synaptic currents, cells were voltage-clamped at −70 mV. A monopolar aCSF-filled electrode placed in the vicinity of the cell (∼20 μM) were used to evoke excitatory postsynaptic currents (EPSCs) 50 ms apart at 0.2 Hz intervals. After a 5-minute baseline, high-frequency stimulation (four 1-s stimulations at 100-Hz applied every 10 s) was delivered, followed by a 10 min recording. Access resistance (<20 MΩ) was assessed every 3 minutes, and recordings were accepted for analysis if changes were <15%.

#### Pharmacological manipulations

Stock solutions of CORT (5 mM; Tocris, UK) and 7,9-Dihydroxiyflavone (DHF) (100 mM; Tocris, UK) dissolved into DMSO were stored at -20°C until used. Stock solutions were diluted in aCSF for a final concentration of 100 nM of CORT and 20 µM of DHF. After incubating for 1 hour, slices were transferred from the drug-containing bath into fresh aCSF and patched within 4 hours.

### Corticosterone immunoassays

Blood (∼15 µL) was collected from the tail vein into ice-cold Microvette® capillary tubes (Sarstedt, Germany) and centrifuged at 8100 RPM for 25 minutes at –4 °C using an Eppendorf 5430 R centrifuge. Serum samples were processed and analyzed using the DetectX® Corticosterone Immunoassay Kit (Arbor Assays, USA). For processing, 5 µL of serum were incubated with 5 µL of dissociation reagent (Arbor Assays) for approximately 8 minutes, followed by dilution with 490 µL of 1X assay buffer (Arbor Assays), resulting in a final 1:100 dilution. Processed samples were stored at – 20 °C until analysis. All samples were analyzed in triplicate on the same day. When possible, repeated samples from the same animal were analyzed on the same assay plate. Plates with standard curves exhibiting an R² < 0.99 were reanalyzed to ensure accuracy. Corticosterone concentrations were reported in ng/mL.

### Brain-derived neurotrophic factor immunoassays

Mice (CRH-IRES-Cre × Ai14 [tdTomato]) were anesthetized with isoflurane, and their brains were rapidly extracted and submerged in ice-cold aCSF for ∼5 minutes. Coronal brain slices (100 µm) containing the PVN were prepared using a vibratome (VT1200 S, Leica) in ice-cold aCSF. Slices were inspected under a fluorescence microscope to identify CRH⁺ neurons expressing tdTomato. PVN regions containing fluorescent signal were microdissected using a scalpel under ice-cold conditions, collected into microcentrifuge tubes, flash-frozen on dry ice, and stored at –80 °C until processing.

For BDNF quantification, samples were homogenized in 500 µL of a custom-made protein extraction buffer (prepared according to kit specifications) using a TissueLyser LT (Qiagen). Homogenates were centrifuged, and the supernatant was aliquoted and stored at –80 °C. BDNF levels were measured using a Mouse BDNF ELISA Kit (EZ0309, Bolster, USA) and quantified using a SpectraMax 190 plate reader (Molecular Devices). All samples were run in triplicate on the same day. Total protein content was assessed using the Pierce BCA Protein Assay Kit (ThermoFisher), and BDNF concentrations were normalized to total protein. Results are expressed as pg of BDNF per mg of protein.

### Optogenetics

Continuous optical stimulation (1 s every 5 s) with blue light (465 nm) reliably activates intracellular TrkB signaling in cells expressing Opto-cytTrkB(E281A)-HA (*50*). For stimulation, a 400 µm core-diameter fiber optic cable (Doric Lenses; MFP_400/430/1100-0.48_2m_FC-MF2.5) was connected to a 465 nm laser, which was controlled by a pulse generator to deliver 1-second light pulses (20 mW) every 5 seconds over a 15-minute period.

### Fiber photometry recordings

Calcium transients in freely moving mice were recorded using a Doric fiber photometry system. The setup consisted of two excitation LEDs (470 nm and 405 nm), controlled by an LED driver and a Doric Studio-compatible console (Doric Lenses). The 470 nm LED (Ca²⁺-dependent signal) was modulated at 208.616 Hz, and the 405 nm LED (isosbestic control) at 572.205 Hz. Signals were demodulated in real time using lock-in amplification. Excitation light was delivered via a Doric Mini Cube filter set (FMC5_E1(405)_E2(460–490)_F1(500–550)_S), coupled to a 400 µm core-diameter fiber optic patch cable (Doric Lenses; MFP_400/430/1100-0.48_2m_FC-MF2.5) connected to the implanted ferrule. LED power at the fiber tip was adjusted to ∼30 μW. Emitted fluorescence was collected through the same fiber and detected by a photoreceiver (Newport Model 2151).

### Fiber photometry data analysis

Fluorescent signal data were acquired at a sampling rate of 100 Hz using the Doric system and exported to MATLAB (MathWorks) for offline analysis with custom-written scripts (https://github.com/leomol/FPA). First, the 470 nm and 405 nm signals were each individually fitted with a second-order polynomial to correct for photobleaching artifacts; the fitted curves were then subtracted from the raw data. Next, a least-squares linear regression was applied to the 405 nm signal to align it with the 470 nm channel. The change in fluorescence (ΔF) was calculated by subtracting the 405 nm Ca²⁺-independent baseline from the 470 nm Ca²⁺-dependent signal at each time point. To minimize the influence of handling- and novelty-induced activity on bleaching correction, a 20-minute baseline recording was obtained in the animals’ home cage (HC). The initial 10 minutes of this baseline were discarded; when the signal reached an asymptotic phase, a 5-minute epoch was selected for curve fitting. Following treatments, subjects were returned to their HC, and Ca²⁺ signals were recorded for an additional 20 minutes. A 5-minute epoch was then selected within the last 10 minutes of this post-treatment recording. Both 5-minute baseline and post-treatment epochs were used to calculate a modified Z-Score (https://github.com/leomol/FPA). Epochs were defined using synchronized video recordings. Compound traces from the selected epochs were presented as continuous time series.

### Stress induction

Electric foot shocks (FS; 0.5 mA) were delivered using either a manually operated scrambler (42.6 × 21 × 29 cm; SMSCK, Kinder Scientific) or an automatic shuttle box (20 × 19.5 × 25.5 cm; Imetronic, France). Acute stress was induced by a single session consisting of ten 3-second foot shocks applied every 27 seconds over a 5-minute period. Stress by immobilization involved restraining animals by tying their limbs and torso onto a flat, highly illuminated (∼630 lux) table using Transpore surgical tape (3M, Germany). Animals remained immobilized for 60 minutes.

### Treadmill training and exercise protocol

The treadmill (TM; LE7808; Panlab, Spain) features a 30 × 10 cm running area with a 10 × 10 cm metal grid at one end. The running area is enclosed by transparent Plexiglas walls and a custom-made lid with an opening to accommodate a photometry optic fiber during experiments. Mice were trained to run on the treadmill more than 24 hours before experiments using the following procedure: subjects were placed on the moving treadmill set at 15 cm/s with a 5% inclination, gently guided forward for 1 minute, and then allowed to run freely for 10 minutes. If a mouse stopped running and was fully dragged onto the metal grid at the end of the treadmill, a mild foot shock (0.1 mA) was delivered. Two training sessions were performed in a single day, with at least 30 minutes of rest between sessions. During the exercise session, mice were placed on the treadmill and allowed to run for 1 hour. Animals were continuously monitored and removed from the session if necessary. Subjects receiving more than 10 seconds of cumulative foot shock during the running session were excluded from the study.

### Behavioral tests

Assessments were conducted between 8:00 and 14:00 hours. Animals’ schedule assignments were balanced across experimental conditions within each daily session and throughout the testing period. Behavioral assessments involved multiple cohorts evaluated at different time points.

#### Dark/Light box

The Dark/Light (D/L) box consists of two rectangular Plexiglas compartments (43 × 21.5 × 30.5 cm; Med Associates) connected by a door. The dark compartment is fully enclosed, painted black, and unilluminated, while the light compartment is open, white, and illuminated (∼240 lux). Animals were habituated to the testing environment by being placed daily in the dark compartment with the door closed for 5 minutes on two consecutive days. Experimental manipulations were conducted more than 2 hours after the final habituation session. On testing day, animals were placed in the dark compartment, and the door was immediately opened to allow access to the light compartment for 5 minutes. All sessions were video recorded (DMK 22AUC03, The Imaging Source), and the following behaviors were analyzed using AnyMaze tracking software (v12, Stoelting): latency to enter the light compartment (seconds), frequency and duration of visits (seconds), and distance traveled (meters) within the light compartment.

#### Contextual Fear Conditioning

Mice were placed on an automatic shuttle box (20 × 19.5 × 25.5 cm; Imetronic, France), where they received ten 3-second foot shocks (0.5 mA) delivered every 27 seconds over a 5-minute period. For contextual fear memory recall, mice were reintroduced to the foot shock chamber for 5 minutes while the duration of freezing, walking, and rearing behaviors were automatically scored (Imetronic, France).

### Statistical analysis

Data was analyzed using GraphPad Prism (v10.2.0) and IBM SPSS Statistics (v29.0.1.1). Group differences were assessed using one-tailed t-tests, one-way, or two-way ANOVAs as appropriate. Bonferroni correction was applied for multiple pairwise comparisons when necessary. Repeated-measures ANOVAs were used to analyze group differences over time. If Mauchly’s test of sphericity indicated violation (P < 0.05), Greenhouse-Geisser correction was applied.

### AI use

Language refinement in the Introduction and Discussion sections were assisted by the large language model ChatGPT (OpenAI, GPT-4, May–September 2025). Some schematics (i.e., mouse icons) were created by the authors using Adobe Illustrator with AI-assisted design tools. All scientific content was conceived, written, and verified by the authors.

### Code availability

Scripts used to analyze fiber photometry are deposited here: https://github.com/leomol/FPA; https://doi.org/10.5281/zenodo.5708470.

## Acknowledgement

We thank Ms. Alexis Passmore, Dr. Leonardo Molina, and Dr. Jianjun Sun for their technical assistance. We are also grateful to Dr. Jonathan Thacker, Dr. Grant Gordon, Sierra Stokes-Heck, Cheryl Breiteneder, Govind Peringod, and Patrick Grouve for their valuable advice and support. We acknowledge the Cumming School of Medicine Optogenetics Core Facility for their continued assistance. Language refinement in the introduction and discussion sections was assisted by the large language model ChatGPT (OpenAI, GPT-4, May–September 2025). Some schematics (i.e., mouse icons) were created by the authors using Adobe Illustrator with AI-assisted design tools. M.R-C. was supported by graduate scholarships from the Cumming School of Medicine and the Killam Trust.

## Author contributions

M.R.C. conceptualized, designed, and conducted the experiments; analyzed the data; prepared the figures; wrote the manuscript; and administered the project. D.B. contributed to experimental design, performed experiments, and analyzed data. S.G.C. assisted in conducting experiments. T.F. and N.D. contributed to conceptualization, experimental design, and supervision of the project. T.F. also contributed to figure preparation and to reviewing and editing the manuscript. M.N.H. contributed to reviewing the manuscript and provided financial support for the project. J.S.B. contributed to conceptualization, experimental design, supervision, and administration of the project. J.S.B. also contributed to reviewing and editing the manuscript and secured funding for the project.

### Data and materials availability

All data are available in the manuscript or the Supplemental information figures.

## Supplemental Information

**Figure S1.**
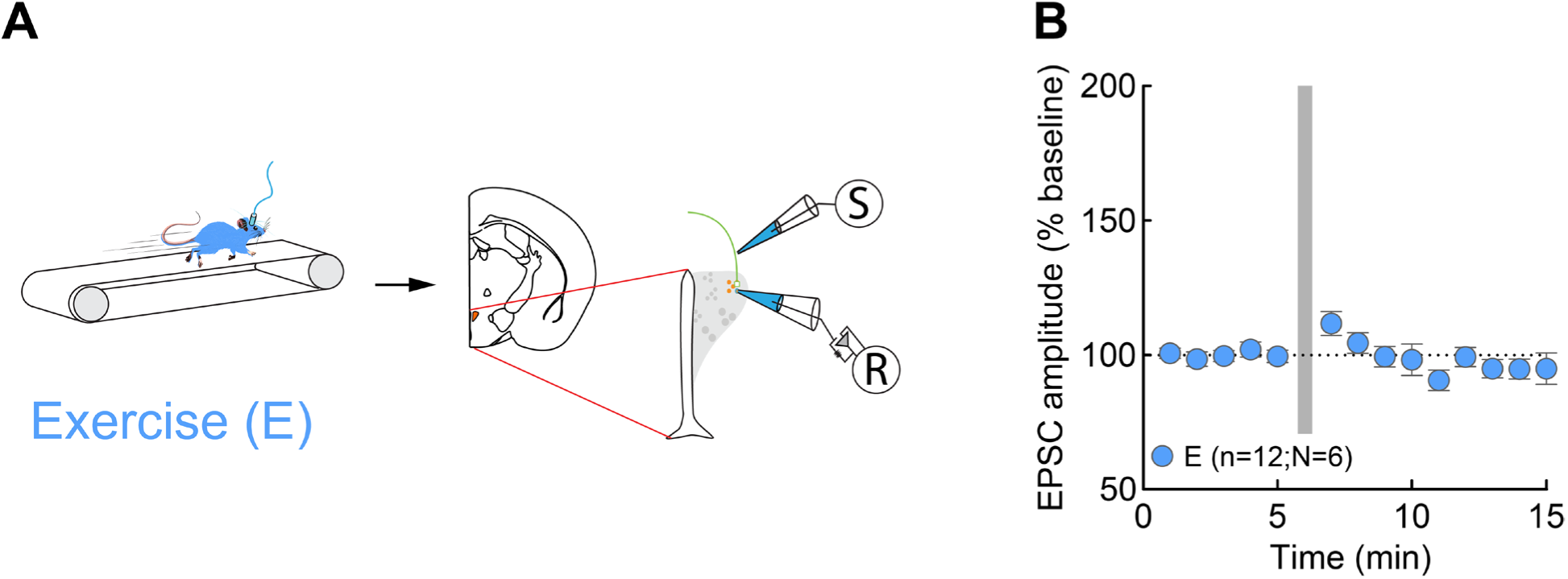
Exercise does not induce STP. **A.** Experimental design for whole-cell patch clamp recordings from CRH^PVN^ neurons of mice exercising on a treadmill for 1 hour. **B**. Summary of excitatory post-synaptic current (EPSC) amplitudes following HFS (grey bar) relative to baseline (BL; doted line). A repeated-measures ANOVA revealed an effect of Time (*F* (13, 143) = 2.175, *P* = 0.013). However, no significant increase in the EPSC amplitudes was observed after HFS (min 5 vs min 7: *P* = 1.00). Bonferroni’s multiple comparisons adjustment was used. Data shown represent the mean ± standard error of the mean (SEM).

**Figure S2.**
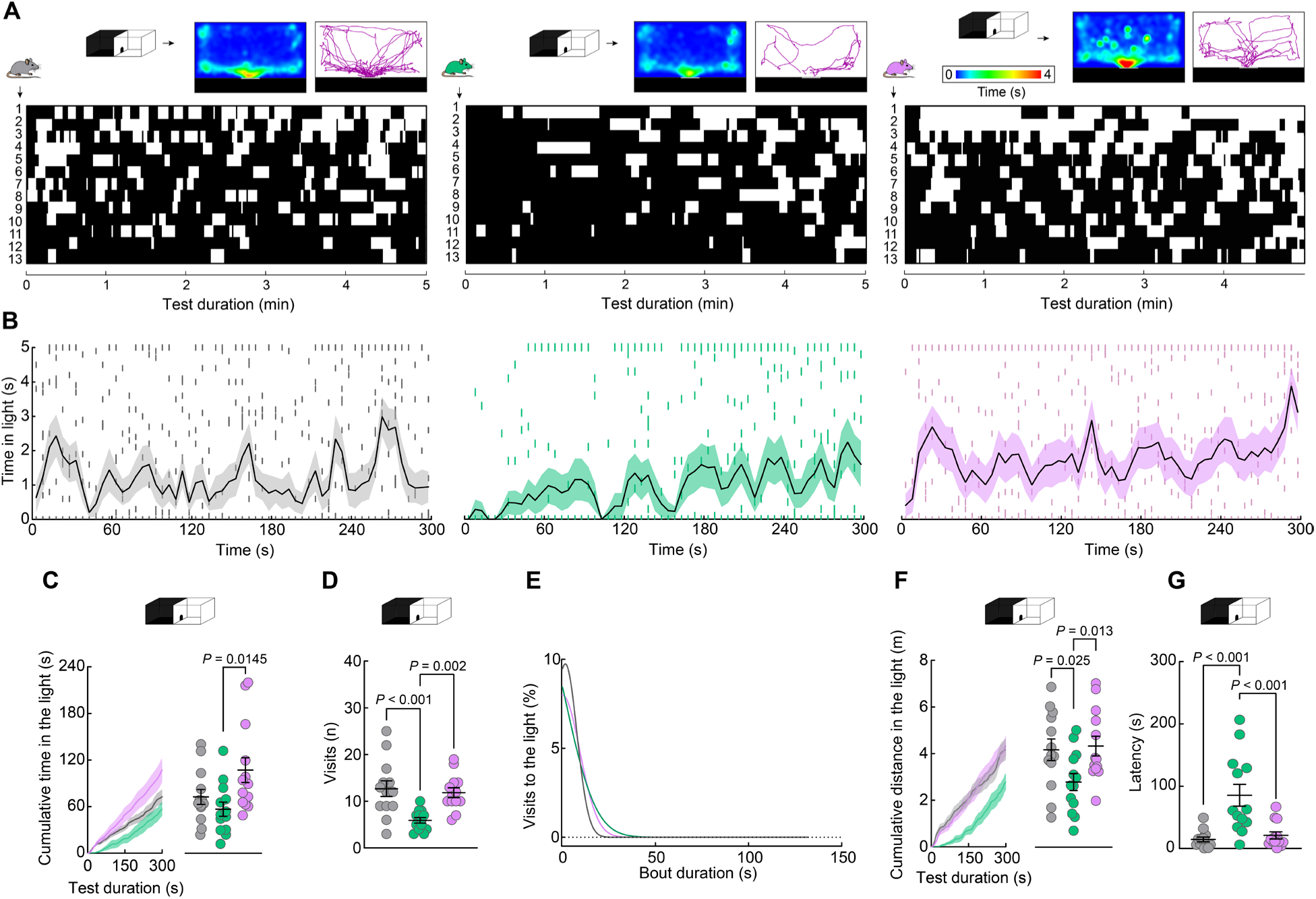
Exercise reverses stress-induced anxiety-like behaviors (five-minute analysis). **A**. Heat maps representing the spatial distribution of the average exploration time in the light compartment (top section of each panel). Tracking plot of the exploration trajectory of a representative animal per group. Behavioral time budget plot showing individual visits to the light (white bars) and dark compartment (black bars) of the DL test during the test (bottom section of each panel). **B**. Frequency analysis of the exploration time in the light compartment binned every five seconds. **C**. Cumulative time in the light compartment. The analysis of the average duration per group revealed S + E mice spent significantly more time in the light compartment compared to the stressed group (*P*= 0.0145; One-way ANOVA: *F* (2, 36) = 4.596, *P* = 0.0167). **D**. Stress significantly reduced the total number of visits to the light compartment compared to N mice (*P* < 0.001), with S + E mice showing increased number of visits compared to S mice (*P*= 0.002; One-way ANOVA: *F* (2, 36) = 9.539, *P* = 0.0005). **E**. No differences in the distribution of bout duration were found between groups. **F**. Cumulative distance traveled in the light compartment over testing minutes. The analysis of the average distance travelled per group revealed that stress significantly reduced animals’ displacement compared with N mice (*P* = 0.025), with S + E mice showing increased levels compared to S subjects (*P* = 0.013; One-way ANOVA: *F* (2, 36) = 4.128, *P* = 0.0243). **G**. Stress also significantly increases the latency to explore the light compartment compared to N mice (P < 0.001), with S + E mice showing a reduced latency compared to S subjects (P < 0.001; One-way ANOVA: *F* (2, 36) = 13.09, *P* < 0.001). Tukey’s multiple comparisons test was used. Data shown represents the mean ± standard error of the mean (SEM).

**Figure S3.**
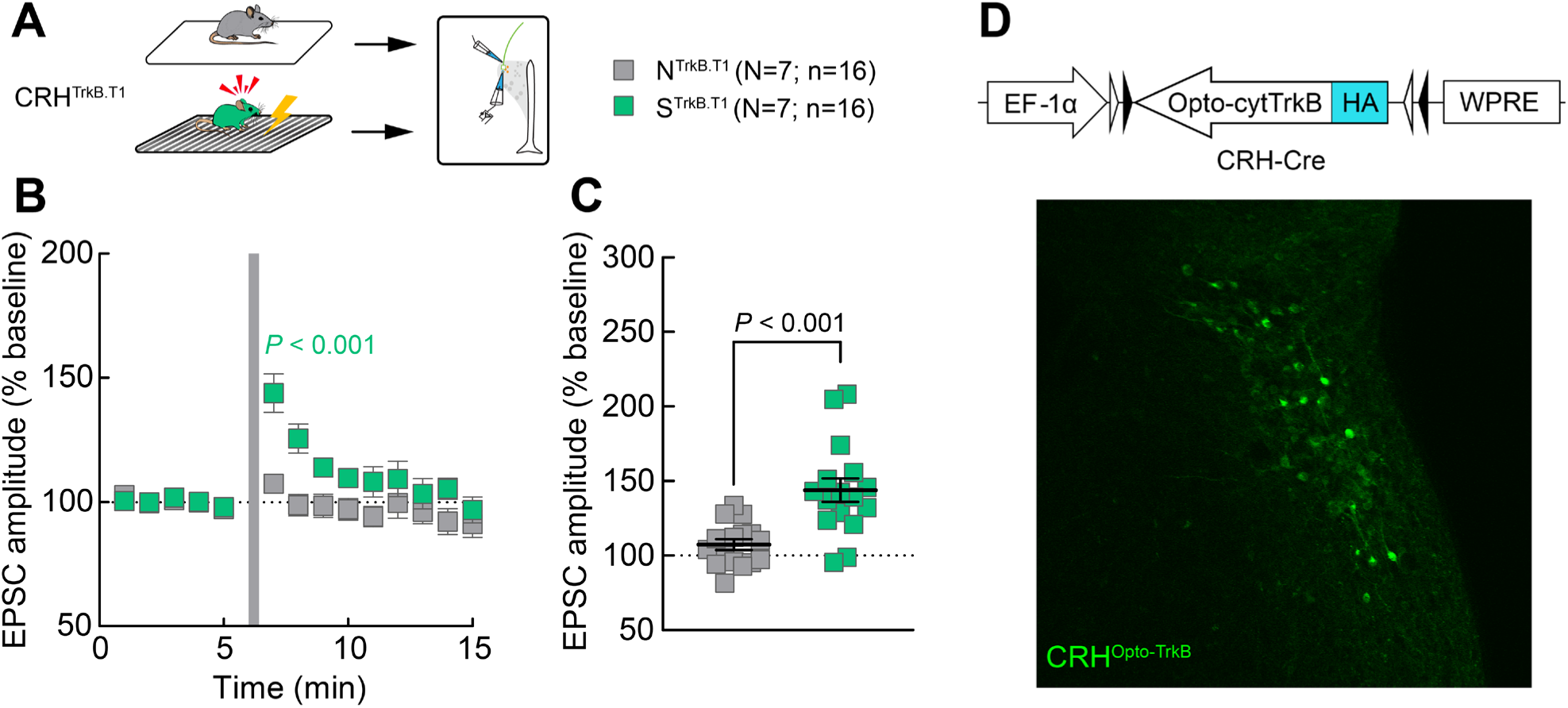
Overexpression of TrkB.T1 in CRH^PVN^ neurons does not induce STP or blocks stress-induced STP. **A.** Experimental design for whole-cell patch clamp recordings from CRH^PVN^ neurons of naïve and stress mice expressing TrkB.T1 receptor in CRH^PVN^ neurons. **B**. Summary of excitatory post-synaptic current (EPSCs) amplitudes following HFS (grey bar) relative to baseline (BL; dotted line). A mixed ANOVA revealed Time (*F* (13, 377) = 8.769, P < 0.001) and Group (*F* (1, 30) = 8.630, *P* = 0.006) main effects, with a Time x Group interaction (*F* (13, 377) = 4.637, *P* < 0.001). EPSC amplitude significantly increased after HFS (min 5 vs min 7: *P* < 0.001) in S^TrkB.T1^ mice only. **C**. Average EPSC amplitude per cell one minute after HFS. S^TrkB.T1^ mice showed higher amplitude compared to N^TrkB.T1^ animals (*P* < 0.001). **D**. Viral construct of the Opto-cytTrkB(E281A)-HA AAV (top). Confirmatory confocal image of CRH^PVN^ cells expressing Opto-cytTrkB(E281A)-HA labeled with Alexa-488 (bottom). Tukey’s multiple comparisons test was used. Geisser-Greenhouse correction was used. Data shown represents the mean ± SEM.

**Figure S4.**
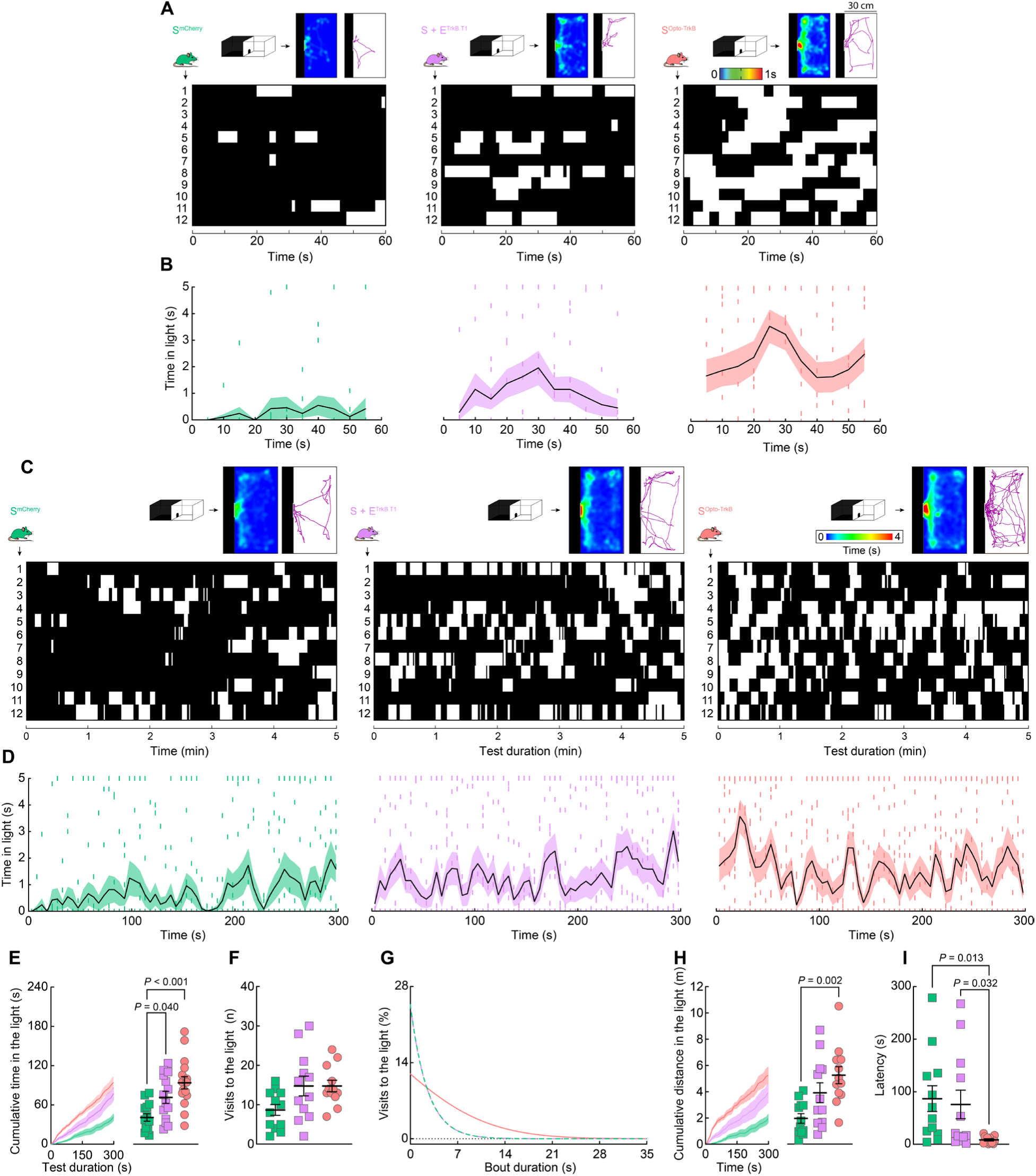
Post-stress activation of TrkB in CRH^PVN^ cells reverses stress-induced STP and anxiety-like behaviors (five-minute analysis). **A**. Heat maps represent the spatial distribution of the average exploration time in the light compartment (top section of each panel) during the first testing minute. Tracking plot of the exploration trajectory of a representative animal per group. Behavioral time budget plot showing individual visits to the light (white bars) and dark compartment (black bars) of the DL test during the first minute of testing (bottom section of each panel). **B**. Frequency analysis of the exploration time in the light compartment binned every five seconds during the first testing minute. **C**. Heat maps represent the spatial distribution of the average exploration time in the light compartment (top section of each panel) during the five minutes of the test. Tracking plot of the exploration trajectory of a representative animal per group. Behavioral time budget plot showing individual visits to the light (white bars) and dark compartment (black bars) of the DL test during the five minutes of the test (bottom section of each panel). **D**. Frequency analysis of the exploration time in the light compartment binned every five seconds during the five minutes of testing. **E**. Cumulative time in the light compartment. The analysis of the average duration per group revealed that both S + E^TrkB.T1^ (P = 0.04) and S^OptoTrkB^ mice (P < 0.001; showed increased time in the light compartment compared to S^mChery^, One-way ANOVA: *F* (2, 33) = 6.584, *P* = 0.004). **F**. No between-group differences were found when analyzing the total number of visits to the light compartment. **G**. No differences in the distribution of bout duration were found between groups. **H**. Cumulative distance travelled in the light compartment. The analysis of the average duration per group revealed that S^OptoTrkB^ mice travelled longer distances in the light compartment compared to S^mChery^ animals (*P* = 0.002; One-way ANOVA: *F* (2, 33) = 7.245, *P* = 0.002). **I**. The analysis of the latency to explore the light compartment revealed shorter latencies in S^OptoTrkB^ mice compared to S^mChery^ (*P* = 0.013) and S + E^TrkB.T1^ subjects (*P* = 0.032; One-way ANOVA: *F* (2, 33) = 4.006, *P* = 0.028). Tukey’s multiple comparisons test was used. Data shown represents the mean ± SEM.

